# Modelling asymmetric count ratios in CRISPR screens to decrease experiment size and improve phenotype detection

**DOI:** 10.1101/699348

**Authors:** Katharina Imkeller, Giulia Ambrosi, Michael Boutros, Wolfgang Huber

**Author notes:** Corresponding authors. (K.I.), (W.H.).

## Abstract

Pooled CRISPR screens are a powerful tool to probe genotype-phenotype relationships at genome-wide scale. However, criteria for optimal design are missing, and it remains unclear how experimental parameters affect results. Here, we report that random decreases in gRNA abundance are more likely than increases due to bottle-neck effects during the cell proliferation phase. Failure to consider this asymmetry leads to loss of detection power. We provide a new statistical test that addresses this problem and improves hit detection at reduced experiment size. The method is implemented in the open source package `gscreend` (submission to Bioconductor pending).

## Background

Genetic perturbation screens are a powerful tool to systematically probe gene function and genotype-phenotype relationships in many different cell types. Their applications include identification of genotype-specific vulnerabilities in human cancer cells^1–3^, discovery of genes involved in drug resistance^5–8^ and virus replication^9,10^.

Currently, the most widespread technology to induce specific genetic perturbations is based on CRISPR (clustered regularly interspaced short palindromic repeats) associated enzymes. In this approach, DNA constructs encoding a guide RNA (gRNA) and the CRISPR associated enzyme are stably integrated into cells, e.g. via lentiviral transduction. The gRNA directs the CRISPR associated enzyme to its sequence-specific target site in the genome. To generate gene knockout perturbations, a common choice of enzyme is the endonuclease Cas9 (CRISPR associated 9)^5,11,12^, which induces DNA cleavage at the genomic site it is directed to. Subsequent DNA repair via non-homologous end joining leads to frame-shift mutations and premature stop-codons, nonsense-mediated RNA decay and finally gene knockout. Alternatively, it is possible to introduce more subtle perturbations such as altered splicing patterns^13,14^ or quantitative modulation of gene expression^15^. To this end, modified CRISPR associated enzymes which function as epigenetic modifiers^16–18^, transcriptional modulators^19–21^ or single-base editors^22,23^ are used.

Pooled screens enable the measurment of the effects of many genetic perturbations in parallel in a single experiment. To this end, a library of gRNAs is introduced into a pool of cells at a low multiplicity of infection such that no more than one gRNA is present in each cell^24,25^. The gRNA sequence simultaneously serves as a barcode that is used to trace which perturbation each cell carries. In the case of negative selection screens, the transduced cell pool is allowed to grow for several divisions during which the relative abundance of cells with a particular gRNA increases or decreases depending on the extent to which the targeted gene determines cell fitness. These effects are detected by amplifying, sequencing and counting the gRNAs before (library or T0) and after (T1) the proliferation phase (Fig. 1A).

**Figure 1:**
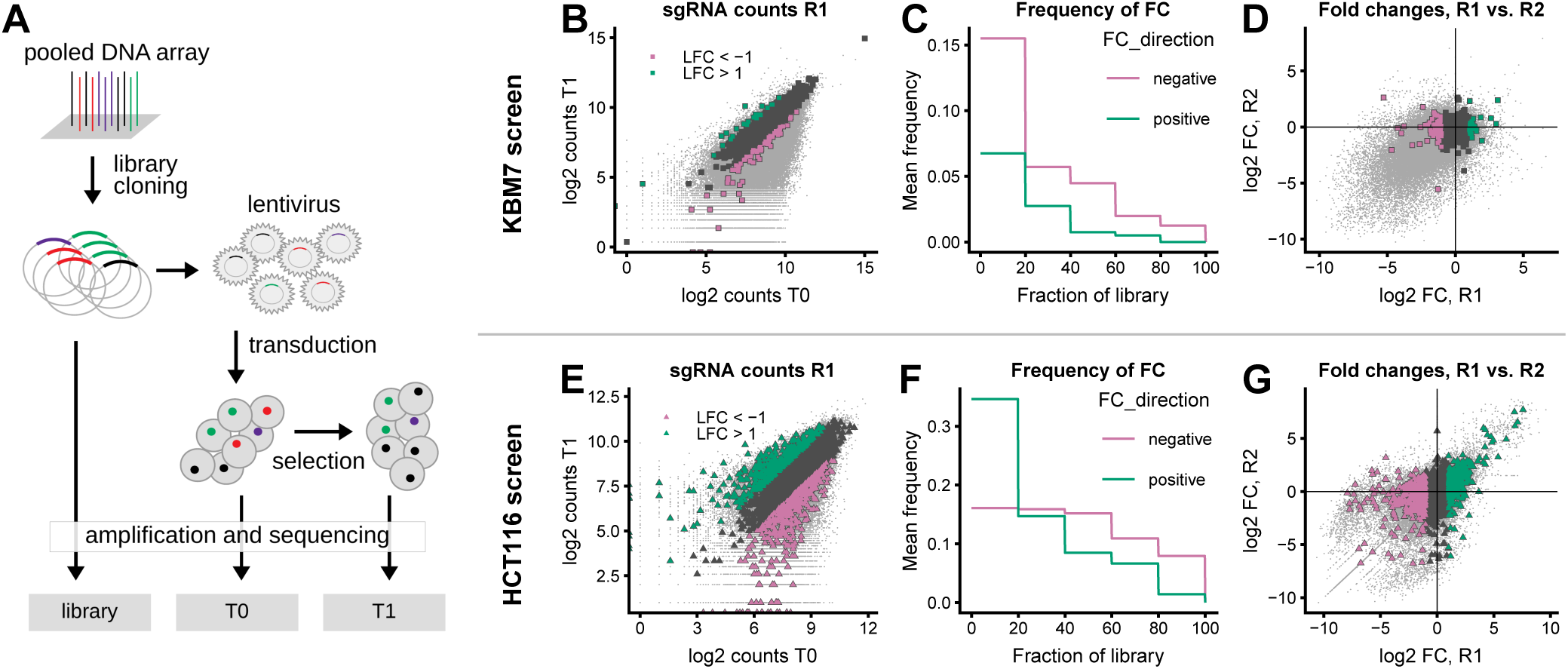
Screening data show an asymmetric distribution of gRNA abundance fold changes. A: Screen setup for measurement of gRNA effect on cell fitness. B-D: KBM7 screen^1^ with highlight on non-targeting controls. E-G: HCT116 screen^2^ with highlight on gRNAs targeting non-essential genes (defined according to Hart et al.^38^). B, E: gRNA abundance at T1 compared to T0 for one of the replicates (R1). Non-targeting control gRNAs and gRNAs targeting non-essential genes (E) are shown by large symbols, all other gRNAs as small grey points. LFC: logarithm of base 2 of inverted before/after ratio calculated as −log2((normalized count at T0 + 1)/(normalized count at T1 + 1)). Colors indicate LFC<-1 (pink) and LFC>+1 (green). C, F: Fraction of non-targeting gRNAs or gRNAs targeting non-essential genes (F) with LFC<-1 (pink) and LFC>+1 (green). The gRNAs were sorted according to their abundance at T0, and the frequencies were calculated within each quintile of the abundance distribution (mean over two replicates). D, G: Comparison of observed LFC for two replicates R1 and R2. Colors correspond to LFC in replicate R1 as in panels B and E.

A typical genome-wide CRISPR screening library for a mammalian genome contains between 70,000 and 120,000 gRNAs^2,5,11,12,26,27^. To ensure statistical power, each gRNA must be represented by a sufficient number of cells during each step of the screen. When designing screening experiments it is convenient to assume that all gRNA are present in the library at approximately the same relative frequencies, and the library composition is summarized by the mean gRNA abundance, also referred to as coverage or representation. This measure is then used to calculate the necessary size of an experiment^24,25^. For a library of 100,000 gRNAs and a desired coverage of 500 for example, 50 million cells (500 times 100,000) must be cultured. Published recommendations on optimal library coverage selection lack precision and range from 200^28^ to 500^24^. Optimizing this choice is a major thrust of this work, since it has substantial consequences on the size, costs and outcomes of an experiment.

To compare the gRNA abundances before and after the proliferation phase, a range of statistical models and computational tools are available^29–35^. Common approaches are to model the joint bivariate null distribution of the normalized counts before and after the proliferation phase, or the null distribution of a univariate summary statistic, the ratio of these counts, hereafter referred to as the “before/after ratio”. gRNAs whose data fall sufficiently outside the null distribution present evidence of a fitness effect of their target gene. Since it is common that each gene is targeted by multiple gRNAs, a subsequent step of the analysis consists of aggregating gRNA-level evidence to the gene level. This can be achieved for example using Bayesian hierarchical modelling^30,34,35^ or robust rank aggregation^29^.

For the null distribution modelling and hypothesis testing, approaches derived from RNA sequencing and differential gene expression analysis have been used^36,37^. However, careful inspection of datasets shows that the distribution of the before/after ratios for negative controls is often asymmetric^34^, which is not the case in RNA sequencing. Such asymmetry means that even in the absence of a fitness effect, a gRNA’s relative abundance is more likely to randomly decrease rather than increase during the screen. Failing to account for such asymmetry (as is done when using the RNA-seq based tools) leads to needlessly elevated false positive rates and/or decreased detection power.

Here, we present a biology-based, generative model that explains the asymmetry of the before/after ratios in pooled CRISPR screens and mechanistically links it to specific steps in the screening experiment. Based on our model, we derive a statistical test that we implemented in the R package `gscreend`, and that enables accurate phenotype detection at reduced experiment size compared to existing approaches. Moreover, through our model we can calculate the minimal experiment size necessary for a given screening library and a required detection power, a point that has never been systematically addressed in any published CRISPR screening protocol.

## Results

### Before/after ratios from pooled genetic screens have an asymmetric null distribution

We studied the distributions of the gRNA counts at T0 and T1 in two pooled CRISPR knockout screens conducted in human cell lines^1,2^ (Fig. 1B-G). After scaling normalization of the counts to the total counts at T0, we computed the logarithm of the ratio of the counts after and before the proliferation phase (logarithmic fold change, LFC, see Methods). We focused on two classes of gRNAs: (a) those that should not have a fitness effect because their sequence does not match any region in the human genome (Fig. 1B-D^1^), and (b) those that target genes that are not essential for cell fitness according to a study by Hart et al.^38^ (Fig. 1E-G^2^). The sign of their LFCs was uncorrelated between replicates, in agreement with the assumption that the LFCs were due to random experimental variability and not due to target-dependent fitness effects (Fig. 1D and G). However, the distribution of LFCs was not symmetric, in particular at the tails: values of LFC<-1 (strongly decreased abundance) were approximately 5-10% more frequent than those with LFC>+1 (strongly increased abundance) (Fig. 1B-C and E-F). These results are qualitatively in accordance with those of Daley et al.^34^.

### Computational simulation of pooled CRISPR screens

To investigate the origin of this asymmetry and possible impact of experimental design parameters, we developed a quantitative model and computational simulation of pooled CRISPR screens. The state space of the model is the tuple of integer counts of the gRNAs, which the model tracks as a function of time throughout the different steps of the screen (Fig. 2A). The temporal evolution of the state is described by endomorphic functions simulating subsampling during transduction, cell splitting and sequencing as well as exponential cell growth according to a gRNA specific growth rate. In our simulations, we considered screens performed with a library of 50000 gRNAs (targeting 12500 genes with 4 independent gRNAs per gene). For 10% of the genes the knockout leads to a growth defect, and for 1% to increased growth. Table 1 summarizes the simulation parameters. A detailed description of the simulation algorithm is provided in the Methods section.

**Table 1:** Parameters of CRISPR screen simulation

| Symbol | Variable in software | Default value | Description |
| --- | --- | --- | --- |
| $C_{\text{virus}}$ | cov_virus | 400 | Coverage during viral transduction. |
| $C_{\text{cells}}$ | cov_cells | 400 | Coverage during cell culture. |
| $C_{\text{PCR}}$ | cov_pcr | 400 | Coverage for PCR amplification. |
| $L$ | lib_width | 7.5 | Library distribution width. |
| $\tau$ | dupl_time | 30 | Cell duplication time in hours. |
| $\phi_{\text{neg}}$ | freq_negfc | 0.1 | Fraction of gRNAs with negative fitness effect. |
| $\phi_{\text{pos}}$ | freq_posfc | 0.01 | Fraction of gRNAs with positive fitness effect. |
| $N_{\text{tot}}$ | n_sgrnas | 50000 | Number of gRNAs in total. |
| $N_{\text{gRNA}}$ | n_sgrnas_per_gene | 4 | Number of gRNAs per gene. |
| $N_{\text{libpcr}}$ | n_repl_lib_pcr | 2 | Number of replicates for library sequencing. |
| $N_{\text{bio}}$ | n_repl_sel | 10 | Number of biological replicates. |
| $N_{\text{biopcr}}$ | n_repl_pcr | 3 | Number of sequencing replicates per biological replicate. |
| $N_{\text{split}}$ | n_splittings | 7 | Number of cell splittings. |
| $\Delta_t$ | - | 72 | Time between cell splittings in hours. |

**Figure 2:**
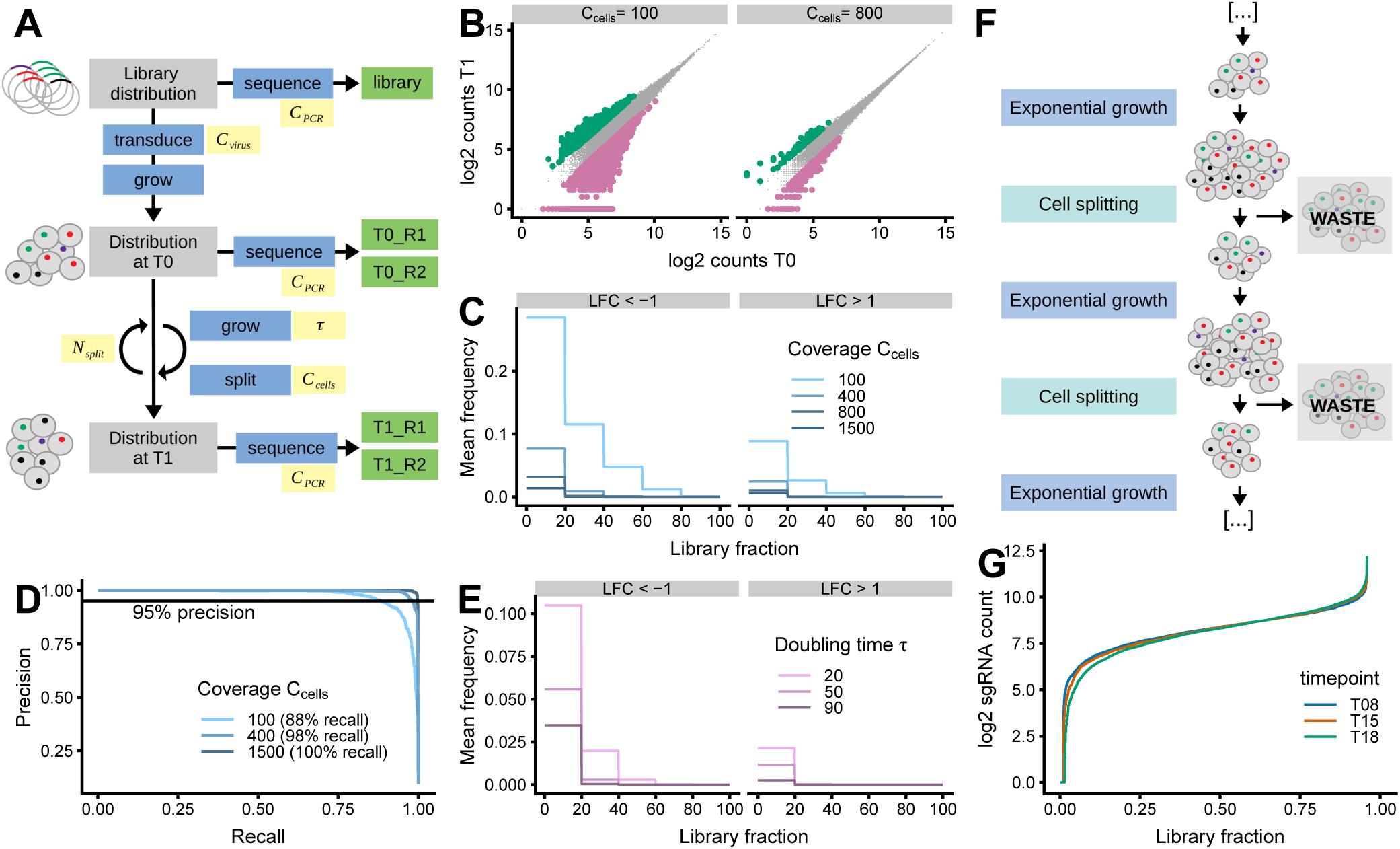
Computational simulation explains how cell splitting causes asymmetry of before/after ratios. A: Schematic representation of the simulation. After generation of an initial gRNA abundance distribution, different functions (blue) are applied to model transduction, cell growth, cell splitting and sequencing. The simulation outputs the gRNA counts obtained by sequencing the plasmid library as well as the cell pools at timepoints T0 and T1 (green, R1 and R2 are technical replicates). The simulation depends on a set of user-defined parameters (yellow, see Table 1). B-E: Simulation results for different values of cell splitting coverage *C*_cells_ and cell doubling time *τ*, while other parameters remain fixed. Only gRNAs without fitness effects are shown. B: gRNA abundance at T1 compared to T0 for simulation with *C*_cells_ of 100 and 800. gRNAs with large observed fold changes are colored (LFC<-1 in pink, LFC>+1 in green). C, E: Fraction of gRNAs with LFC<-1 (left) and LFC>+1 (right) for *C*_cells_ ranging from 100 to 1500 (C) and *τ* ranging from 20 to 90 hours (E). Mean over 5 simulations is depicted. D: MAGeCK-RRA precision-recall curves on data simulated using different values for *C*_cells_ (100, 400 and 1500). The recall at 95% precision is indicated. F: Schematic representation of cell splitting during the proliferation phase of screen, which consists of multiple rounds of exponential growth and random sampling. G: Count distribution of gRNAs targeting non-essential genes at T08, T15 and T18 of the screen in HCT116 cells^2^. gRNAs were ranked according to their abundance and the resulting ranks normalized to [0;1] (library fraction, x axis). On the y axis the counts per gRNA are shown.

### Plasmid library is a better reference sample than T0 cell pool

We first investigated the effect of the choice of reference sample. Previous publications used gRNA counts from either the plasmid library or the T0 cell pool as reference (Fig. 1A)^1,2,27,39^, and it is unclear to what extent this choice influences the analysis outcome. Timepoint T0 is after the antibiotics selection of cells that were successfully transduced, in other words, up to four cell doublings after transduction. Such selection is necessary because at typical multiplicities of infection, only a fraction of cells is infected. In our simulations, we observed that counts of gRNAs targeting essential genes were already decreased at T0, especially for fast growing cells (Fig. S1A). To confirm this experimentally, we transduced pools of Cas9 expressing HCT116 and RKO cells with a genome-wide CRISPR library and selected the successfully transduced cells for 4 days, similar to the period before T0 in a screen. We sequenced the gRNAs in the plasmid library and at T0 and compared their normalized abundances. Similar to the prediction from the simulation, gRNAs targeting essential genes^38^ had reduced abundance at T0 (Fig. S1B). This result implies that plasmid library rather than T0 sequencing should be used as a reference to avoid premature under-representation of gRNAs targeting essential genes.

### The asymmetry of before/after ratios is caused by cell splitting during the proliferation phase

We next investigated the effect of experimental parameters on the distribution of before/after ratios for gRNAs without effect on cell fitness (Fig. 2B-E). The key parameter for cell culture during a screen is the mean gRNA coverage, which is reflected in the number of cells that are seeded after every round of splitting. We found that the smaller the coverage during cell splitting, the greater the asymmetry of before/after ratios (Fig. 2B-C). Higher levels of asymmetry lead to impairment of phenotype detection by MAGeCK-RRA^29^, a current state-of-the-art analysis tool, which lost 10% of recall at 95% precision when, e.g., reducing the cell splitting coverage from 400 to 100 (Fig. 2D). Similarly, the asymmetry increased when using faster growing cell lines, as we observed in our simulations (Fig. 2E) and in experimental datasets (Fig. S2)^39,40^. We also tested the effect of other parameters, such as coverage during transduction or PCR. These, however, only marginally influenced the asymmetry of before/after ratios in our simulations (Fig. S3). Decreasing the cell splitting coverage led to up to 20% of gRNAs with LFC<-1, whereas for similar changes in PCR or transduction coverage, this fraction was only 3% (Fig. 2C and S3).

We conclude that transduction and PCR do not cause major technical biases in the data and that it is better to sequence the gRNA pool in the plasmid library rather than at T0. The observed asymmetry however can be mechanistically explained by multiple rounds of cell splitting bottlenecks and exponential growth (Fig. 2F). With every round of exponential growth followed by random sampling of cells, the distribution of gRNA abundances gets wider, i.e. there are more and more gRNAs that are underrepresented in comparison to the mean gRNA coverage. We confirmed the gradual broadening of the abundance distribution of gRNA targeting non-essential genes^38^ in a published dataset from a CRISPR-knockout screen performed in HCT116 cells (Fig. 2G)^2^.

### Wide initial gRNA abundance distributions increase asymmetry of before/after ratios

Since the observed asymmetry is caused by broadening of the gRNA abundance distribution, we hypothesized that the width of the gRNA abundance distribution in the plasmid library also influences the data quality of the screen. A measure of the width of this distribution, i.e. the difference in abundance between low and high abundant gRNAs, is the ratio between the 90% and 10% percentiles. This measure, elsewhere also named “skew ratio”^24^, will hereafter be referred to as “distribution width”. If, for example, the most abundant 10% gRNAs of a library have an abundance higher than 500 whereas the least abundant 10% have less than 100 counts, the distribution width is 5.

We performed simulations starting from three different gRNA libraries with varying distribution width (Fig. 3A). Our simulations showed that with higher width the reproducibility between replicates decreased (Fig. 3B) and at the same time the frequency of gRNAs with LFC<-1 increased (Fig. 3C). Experimental data from a screen conducted using two plasmid libraries with different distribution widths confirmed this finding (Fig. 3D-E)^27^. Using plasmid libraries with narrower gRNA abundance distributions thus increases data quality by reducing the asymmetry of the distribution of before/after ratios.

**Figure 3:**
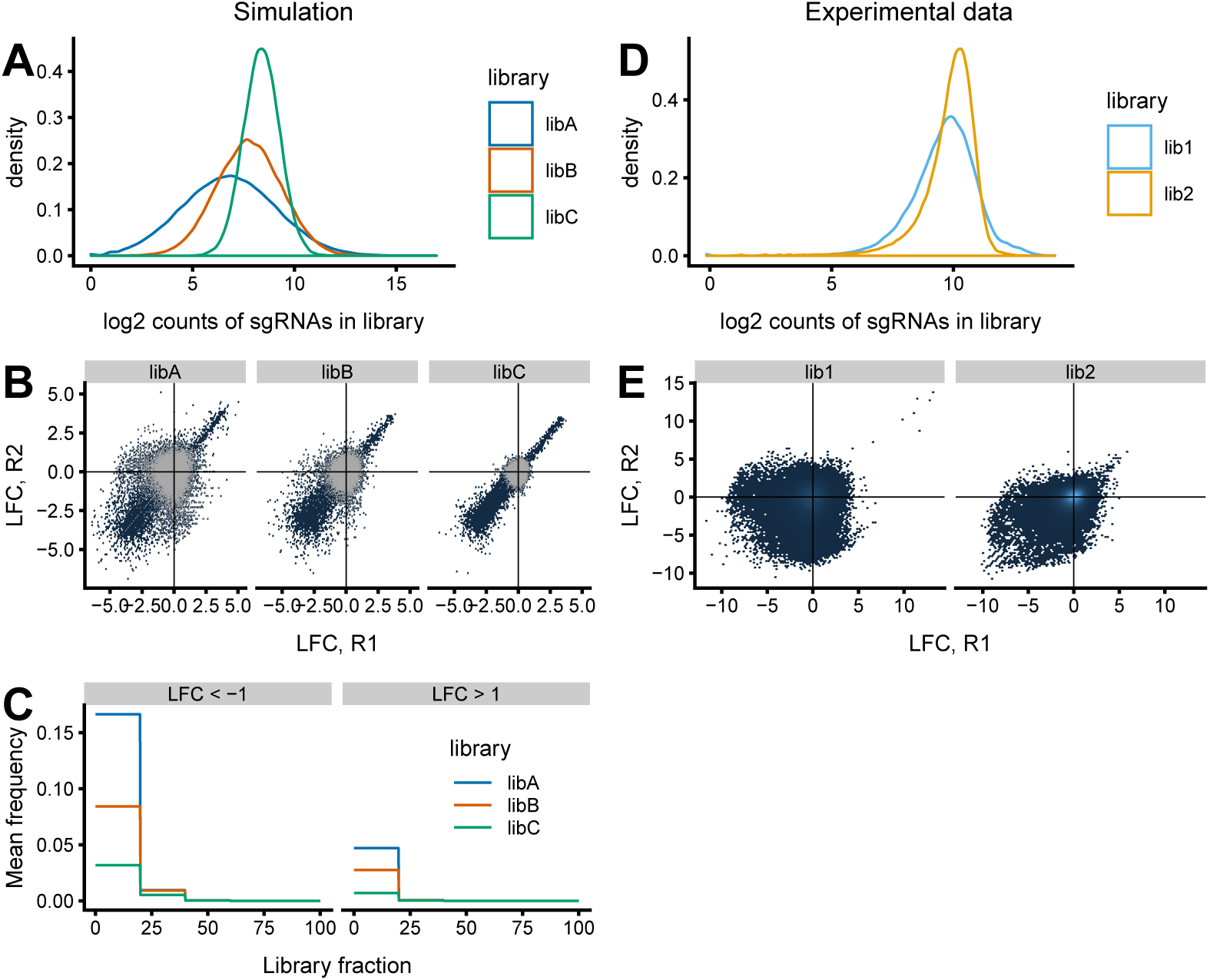
Wider gRNA abundance distributions in the plasmid library increase asymmetry of before/after ratios. A-C: Simulation results using three libraries with different widths of gRNA abundance distribution. A: log2 count distribution of simulated libraries libA, libB and libC. Distributions were generated as log-normal distributions with same log-mean, but differing log-sd and then size-normalized. Library distribution widths: 66.5 (libA), 17.8 (libB), 4.8 (libC). B: Reproducibility of LFC between two replicates. The simulations were performed with libA, libB or libC. All gRNAs are shown in blue, gRNAs without fitness effect are highlighted in grey. C: Fraction of gRNAs with LFC<-1 (left) and LFC>+1 (right) for simulations with libA, libB or libC. gRNAs used for frequency calculation do not have fitness effects. D-E: Dataset from screen performed in mESCs using two different libraries lib1 and lib2 (libraries V1 and V2 respectively from Tzelepis et al.^27^). Library distribution widths: 8.8 (lib1), 5.0 (lib2). D: log2 count distributions of lib1 and 2, normalized to total count. E: Reproducibility of LFC between two replicates in the screens performed with lib1 or lib2. All gRNAs are shown in blue.

Furthermore, we also found that the gRNA sequence composition of a library correlates with its width and that gRNAs with specific sequence properties are more likely to be over- or underrepresented (Fig. S4). To show this we selected five datasets from published CRISPR libraries^1,27,41,42^. These libraries have different distribution widths ranging from 2.4 to 8.8 (Fig. S4A-B). To examine the sequence composition we generated probability sequence motifs for the least and most abundant gRNAs (Fig. S4C-D)^43^. Wider libraries tend to have poly-G-stretches in low abundant gRNAs and many T nucleotides in high abundant gRNAs, especially in the second half of the sequence (Fig. S4E). This is probably due to sequence specific biases during the generation of the plasmid library, for example during synthesis or PCR amplification of gRNAs.

### New statistical method for improved phenotype detection

We showed that before/after ratio distributions in pooled CRISPR screens are asymmetric due to technical artifacts arising during the cell proliferation phase. This asymmetry is influenced not only by cell splitting parameters but also by the width of the gRNA abundance distribution in the plasmid library. In principle, it would be possible to eliminate this asymmetry by using plasmid libraries with minimal distribution width and to perform the screen at very high coverage. However, since this is generally neither feasible nor economically reasonable, we developed a new statistical test that accounts for the asymmetric null distribution. The underlying idea of our method is to use a skew normal distribution to model the LFC null distribution.

The workflow of our new analysis method `gscreend` is depicted in Figure 4A. After scaling of gRNA counts and calculation of LFCs, the data is split into slices according the gRNA abundance in the reference sample (e.g. plasmid library). We introduce this stratification since it allows the parameters of the null distribution to be different for gRNAs with low and high abundance, consistent with what we observed in datasets. We model the LFCs in each stratum as a mixture of a parametric null distribution, the skew normal, and an unspecified alternative distribution^44,45^. The first mixture component corresponds to gRNAs without fitness effect, the second to those with effect, where we assume that these are only a minority. `gscreend` uses least quantile of squares regression^46^ to fit the null distribution to the LFCs in each stratum. Least quantile of squares regression fits a model by only taking into account a defined proportion of residuals, e.g., those between the 10% and 90% percentiles. In contrast to the commonly used least sum of squares regression, it is thus more robust to outliers. In the `gscreend` workflow the resulting null-models for every stratum are used to calculate p-values, which are then employed to rank the gRNAs. Subsequently, robust rank aggregation^29,47^ is applied to aggregate the ranked gRNA list to the gene level.

**Figure 4:**
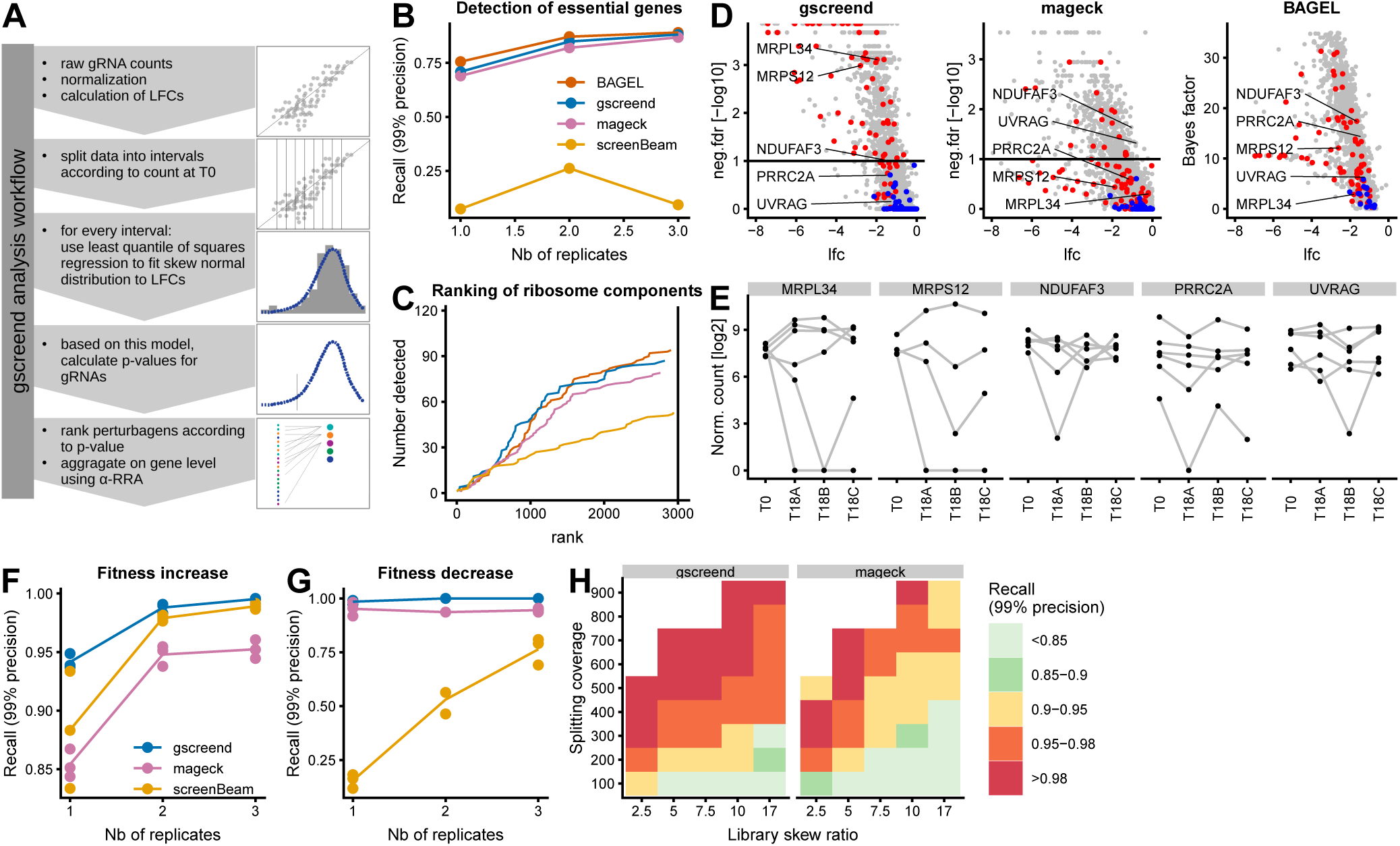
gscreend increases phenotype detection accuracy. A: `gscreend` analysis workflow. B-E: Comparison of gene ranking by BAGEL, MAGeCK, screenBeam and `gscreend` for CRISPR knockout screen performed in HCT116 cells^2^. B: Recall at 99% precision (as in Fig. 1D) for analysis of 1 to 3 biological replicates of timepoint T18. Essential and non-essential genes were defined according to Hart et al.^38^. C: Ranking of genes encoding ribosome components (GO: structural constituent of ribosome) by the four different analysis tools and using all 3 biological replicates. D: Volcano plots illustrating analysis results for genes encoding ribosomal components (red) and non-essential genes^38^ (blue). Horizontal lines indicate FDR thresholds of 1%. E: Log2 normalized abundances of gRNAs targeting the five selected genes at timepoint T0 and three replicates of timepoint T18. F-G: Recall at 99% precision by MAGeCK, `gscreend` and screenBeam for simulated data of 1 to 3 biological replicates. Other simulation parameters: library distribution width 7.5, cell splitting coverage 200, doubling time 30h. Precision-recall curves were calculated for detection of essential (F) and growth-suppressing (G) genes. H: Recall at 99% precision of essential genes for simulations with different library width and cell splitting coverage. Genes were ranked using `gscreend` (left) or MAGeCK (right).

We first tested how well `gscreend` performed in accurately ranking genes in experimental datasets. Using a published list of essential and non-essential genes^38^ we calculated the recall at 99% precision (as in Fig. 2D) of our and other tools^29,30,32^. `gscreend` outperformed MAGeCK and screenBeam when ranking genes in a CRISPR-knockout screen performed in HCT116 cells (Fig. 4B)^2^. BAGEL was the only tool that had a better precision-recall performance than `gscreend` on these data. However, its algorithm was trained on the same benchmark set of essential and non-essential genes that we used here to calculate precision-recall statistics, which might explain some of this performance. Indeed when ranking components of the ribosome, whose knockout is likely to be lethal, `gscreend` outperformed BAGEL, MAGeCK and screenBeam, especially within the 1000 lowest ranked genes (Fig. 4C). In order to illustrate some examples, we highlighted the results for five selected genes (Fig. 4D+E). *MRPL34* and *MRPS12* (components of the mitochondrial ribosome) are detected with low rank only by `gscreend`, although their gRNA abundance profile indicates that they are truly essential, because two of the corresponding gRNAs are strongly depleted at T18 in all three replicates (Fig. 4E). The other three genes *NDUFAF3, PRRC2A* and *UVRAG* are assigned low ranks in BAGEL or MAGeCK but remain above the 1% false discovery rate (FDR) threshold in the `gscreend` results (Fig. 4D). Their gRNA abundance profile indicates that the observed negative LFCs are technical artifacts, because they were not reproduced between replicates (Fig. 4E). Taken together, these results indicate that `gscreend` delivers superior accuracy in ranking and identifying essential genes in pooled negative selection screens.

We also investigated whether our method is robust against different levels of the asymmetry of before/after ratios by simulations. Similar to what we found using the experimental data, `gscreend` had a better ranking accuracy than the other tools when detecting genes that either increase or decrease cell fitness(Fig. 4F+G). When increasing the asymmetry by reducing cell splitting coverage or increasing library distribution width, the method maintained a better ranking accuracy than MAGeCK-RRA (Fig. 4H). `gscreend` enables reduction of the cell splitting coverage by approximately 50% for a library distribution width of 7.5: using 300 (`gscreend`) instead of 600 (MAGeCK) mean gRNA coverage maintained at least 95% recall at 99% precision (Fig. 4H). For libraries with larger distribution widths the gain in accuracy is even more substantial.

### Implications for the design of screening experiments

Based on the findings reported above, we suggest that when designing a screen, the distribution width of the gRNA plasmid library should first be measured. Based on this measure, our simulation tool (Fig. 4D, left panel), can then be used to predict the corresponding optimal coverage to perform cell culture (summarized in Table 2). For PCR and transduction, since these only mildly affect data quality, the respective coverages should be chosen in the same range as the cell splitting coverage.

**Table 2:** Recommended screening coverage for different library distributions

| library width | screening coverage |
| --- | --- |
| 2.5 | 200 |
| 5 | 300 |
| 7.5 | 300 |
| 10 | 400 |
| 17 | 400 |

## Discussion

Accurately detecting phenotypes in pooled genetic perturbation screens is key to generating hypotheses that justify follow-up. Screens that correctly distinguish all genes that negatively or positively regulate cell fitness can be used not only to identify the strongest ‘hits’, but also to measure subtle differences in growth rate and thus map whole pathways and potentially identify mechanisms.

High data quality and accurate analysis is first of all achieved by understanding how the experimental design influences the results. Simulations of CRISPR-based screens have been published^48^, however, our study is the first one to systematically explore the influence of experimental design on phenotype detection in pooled screens. We show that gRNA coverage during PCR and transduction, providing this is chosen in the same range as the cell splitting coverage, only marginally influences data quality and that screens are best analyzed when using plasmid library sequencing as reference. We do not discuss the influence of the multiplicity of infection during viral transduction, as there is already literature and a good model available to address this point^25^. Our most important novel finding is that the asymmetry of the distribution of before/after ratios is caused during the proliferation phase of pooled negative selection screens. Multiple consecutive rounds of cell splitting and exponential growth gradually lead to random loss of low abundant gRNAs.

Using this knowledge of the asymmetric null distribution of before/after ratios, we developed a new statistical test that improves phenotype detection. `gscreend` outperforms existing analysis methods, which assume that the null distribution is symmetric. From the point of view of screen design, our method enables reduction of experiment size by ca. 50% compared to other tools, because it maintains high analysis accuracy throughout a broad range of experimental settings. Especially for experiments that are limited by their size, because for example they use primary cell cultures^37,49^, our tool may help to improve phenotype detection.

Our results also provide indications on how to optimize the experimental design by choosing the screening coverage according to the width of the utilized plasmid library (Table 2). Intriguingly, the width of the library distribution is the limiting parameter that dictates the minimal size of a screening experiment. It would thus be possible to strongly reduce the experiment size by using a library with a narrower distribution. Our analyses indicate that gRNAs with specific sequence characteristics are likely to be over- or underrepresented in gRNA libraries obtained using arrayed synthesis approaches and cloning. We hypothesize that the broadening of library distribution is due to sequence specific differences in synthesis or amplification efficiency. A recently published approach to synthesize covalently-closed-circular-synthesized (3Cs) gRNA libraries may thus be a promising technology for substantial reduction of library width and experiment size^50^.

Finally, the discovery of sequence specific representation differences of gRNAs in a library also has important implications for the evaluation of their gene knockout efficiency. gRNAs with specific sequence properties might seem more efficient than others simply because they are less abundant in the library and thus more likely to suffer from the here described asymmetric loss phenomenon^27,51^.

## Conclusion

We conclude that the asymmetry of the before/after ratio distribution in pooled CRISPR screens is primarily caused by insufficient coverage of gRNAs during the cellular growth phase of a screen. Our results can be used to predict necessary experiment sizes, which are most importantly dictated by the width of the plasmid library. Our new R package `gscreend` takes into account the asymmetry of the null distribution and thus improves phenotype detection at reduced experiment size.

## Methods

### Experimental datasets

The following datasets from published CRISPR knockout screens were used: screen in KBM7 cells^1^ (Fig. 1); screen with TKO library in HCT116 cells, timepoints T08, T15, T18, raw data file version 1^2^ (Fig. 1, 2, 4); screen in mouse ESC using mouse genome wide libraries V1 and V2^27^ (Fig. 3). gRNA counts data from the DepMap project^39,40^ was downloaded together with a dataset of cell doubling times^3^ (Fig. S2).

Data from library and T0 sequencing used in Fig. S1 was collected during a CRISPR screen in HCT116 and RKO cells. The 90k Toronto human Knockout pooled library (TKO) was a gift from Dr. Jason Moffat (#1000000069, Addgene). Plasmid library was amplified using ElectroMAXTM Stbl4TM cells (Invitrogen) accordingly to the manufacturer’s protocol. Library vector was transfected into HEK293T cells (ATCC) with TransIT-LT1 (Mirus Bio) transfection reagent along with psPAX2 (#12260, Addgene) and pMD2.G (#12259, Addgene) packaging plasmids to produce lentivirus. HCT116 and RKO cells (ATCC) stably expressing Cas9 (#73310, Addgene) were infected in presence of 8 *µ*g/ml polybrene (Merck Millipore) with the 90k TKO gRNAs library at a MOI equal to 0.3 such that each gRNA was present in 500 cells on average. The day after, puromycin-containing medium was added to the infected cells for 48 hours. On day four after transduction, a portion of cells were harvested as T0 timepoint. Genomic DNA from cell pellets was extracted using QIAamp DNA Blood Maxi kit (Qiagen). To amplify the gRNA sequences a total of 140 PCR reactions were performed using 1*µ*g of genomic or plasmid library DNA each (250 fold coverage), Q5 Hot Start HF polymerase (NEB), and primers harboring the Illumina TruSeq adapter sequences. PCR products were purified using DNA Clean & Concentrator TM-100 (Zymo Research) and MagSi-NGSprep Plus beads (Steinbrenner). Sample concentrations were measured using Qubit HS DNA Assay (Thermo Fisher). Library amplicon size was verified using DNA High Sensitivity Assay on a BioAnalyzer 2100 (Agilent) and then sequenced on a NextSeq (Illumina) by 75bp single-end sequencing and addition of 25% PhiX control v3 (Illumina). gRNAs were counted using the count function with automatic sequence trimming provided by MAGeCK^29^.

### Simulation of pooled CRISPR screens

We simulate a complete pooled CRISPR-knockout screen, providing output files that represent gRNA counts after sequencing of the plasmid library and T0 and T1 samples (see also Fig. 2A). The simulation depends on several parameters that reflect the experimental setup (see Table 2).

In a first step, the abundance *n*_lib,*g*_ of every gRNA *g* (where *g* = 1, *…, N*_tot_) in the plasmid library is sampled from a lognormal distribution *LN* (*µ, σ*), where *µ* = 5 and *σ* is chosen to match the user-specified library distribution width *L*. We chose *µ* = 5 because resulting distributions resemble those seen in experimental data. The sequencing counts 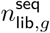 are obtained by making *C*_PCR_*N*_tot_ draws from the multivariate hypergeometric distribution with probabilities 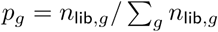. This is repeated *N*_libpcr_ times, to model the technical replicates.

In the next step the abundance of gRNAs in the transduced cell pool *n*_trans,*g*_ are obtained by making *C*_PCR_*N*_tot_ draws from the multivariate hypergeometric distribution with probabilities 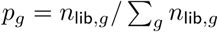.

The pool of gRNAs of total size *N*_tot_ is partitioned into three sets: gRNAs without effect on cell fitness (Γ_neutral_), gRNAs increasing cell fitness (Γ_pos_) and gRNAs decreasing cell fitness (Γ_neg_>). The sets Γ_neutral_, Γ_pos_ and Γ_neg_> have respective sizes *N*_neutral_, *N*_pos_ and *N*_neg_> such that *N*_neg_> = *ϕ*_neg_>*N*_tot_, *N*_pos_ = *ϕ*_pos_*N*_tot_ and *N*_neg_> + *N*_pos_ + *N*_neutal_ = *N*_tot_. gRNAs from the different categories are assigned to essential, non-essential or growth-supressing genes according to *N*_gRNA_.

In general, the cell proliferation-induced change in abundance of gRNA *g* between times *t* and *t* + Δ_*t*_ can be modeled as *n*_*g*_(*t* + Δ_*t*_) = *e*^*β*^*n*_*g*_(*t*), where *β* is the baseline cellular growth factor between two splittings and Δ_*t*_ the time between two splittings. *β* for a specific cell doubling time *τ* can thus be calculated as 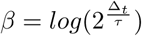.

A gRNA specific growth rate *β*_*g*_ is then derived from *β*_baseline_ such that:

*β*_*g*_ = *β* for every *g* ∈ Γ_neutral_,
*β*_*g*_ = *β*(1 + *ϵ*) for every *g* ∈ Γ_pos_,
*β*_*g*_ = *β*(1 − *ϵ*) for every *g* ∈ Γ_neg_,

where *ϵ* is randomly chosen from {0, .01, 0.02, …, 0.2}.

The gRNA abundances at time t0 are calculated from the abundances in the transduced cell pool as:

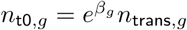 (real numbers are converted to integer by only taking the integer part).

The sequencing counts from T0 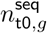 are obtained by making *C*_PCR_*N*_tot_ draws from the multivariate hypergeometric distribution with probabilities 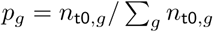. This is repeated *N*_biopcr_ times, to model the technical replicates.

Next the proliferation phase of the screen is simulated *N*_bio_ independent times to model the biological replicates. For i = 1 *…* N_split_, the gRNAs abundances after cell splitting *n*_i,split,*g*_ are obtained by making *C*_cells_*N*_tot_ draws from the multivariate hypergeometric distribution with probabilities *p*_*g*_ = *n*_i,*g*_*/Σ*_*g*_ *n*_i,*g*_. This random sampling step is followed by an exponential growth step 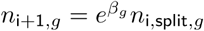. After completion of all cell splittings the gRNAs in all biological replicates (timepoint T1) are sequenced by making *C*_PCR_*N*_tot_ draws from the multivariate hypergeometric distribution with probabilities 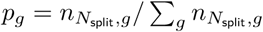. This is repeated *N*_biopcr_ times, to model the technical replicates.

For the analyses shown in Figures 2B-E, 3A-C and 4F-H, *C*_PCR_ and *C*_virus_ are chosen as indicated in the following table. The values of *C*_PCR_ and *C*_virus_ are chosen so that the 10% percentile of low abundant gRNAs in the library have a coverage of 100 fold.

| library width | $C_{\text{PCR}}$ and $C_{\text{virus}}$ |
| --- | --- |
| 2.5 | 120 |
| 5 | 250 |
| 7.5 | 380 |
| 10 | 500 |
| 17 | 850 |

### Normalization and LFC calculation

Counts from experimental data were normalized using size normalization to the total read counts of the reference sample. This was not necessary for simulated datasets, because these already had the same read counts. For a given gRNA with count *n*_*lib*_ in the library and *n*_1_ at timepoint T1, the log fold change was calculated as 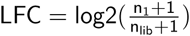. Pseudo-counts had to be added for division and log-transformation since some of the low abundant gRNAs had 0 counts in one or more of the replicates.

### Library width calculation

The width of a distribution of gRNA abundances can be quantified by calculating the ratio between the 90% and 10% percentile of the distribution^24^: 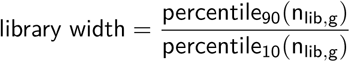. *n*_lib,g_ is the distribution of gRNA abundances in the library.

### gRNA sequence composition

The sequence logos in Fig. S4 were generated using the plogo online tool^43^. gRNAs were ranked according to their abundance and the sequence logos generated for the lower and upper 1% of gRNAs.

### gscreend method

`gscreend` is designed to account for asymmetric distribution of before/after ratios in pooled genetic perturbation screens (see also Fig. 4A).

`gscreend` takes (non-normalized) gRNA counts from several samples as its input. One of these is the reference sample (e.g. the library or T0), the others are one or several replicates of a post-screen timepoint (e.g. T1). The counts are scaled (a.k.a. normalized) to the total counts of the reference sample. Log2 fold changes are calculated as described above. The data are split into slices according to the gRNA abundance in the reference sample; the current implementation uses 10 slices split at the 10%, 20%, … quantiles. We use this stratification because the null distributions of the fold changes depend on it, and are fit separately in each stratum.

We model the overall LFC data as a mixture of a parametric null distribution, the skew normal, and an unspecified alternative distribution^44,45^. The first mixture component corresponds to gRNAs without fitness effect, the second to those with effect, and we will assume that these are only a minority. We use the R package fGarch for computations involving the skew normal distribution and use least quantile of squares regression^46^ on a 10%-90% percentile of the log-likelihood to infer the model parameters from the distribution of LFCs (function lbfgs from R package nloptr).

In the next step, for every stratum p-values are calculated for every gRNA. The gRNAs are ranked based on their p-values (if there are multiple replicates, each gRNA gets as many ranks). On this ranking, `gscreend` uses an *α*-RRA algorithm with an *α* cutoff of 5% to aggregate the data to the gene level^29,47^. Gene level LFCs are calculated by averaging the LFCs over all gRNAs belonging to the gene.

### Comparison of analysis tools

Results from the analysis of simulated and experimental data using the different analysis tools were compared as follows:

- `gscreend` analysis was performed with 10%-90% percentile for least quantile of squares method and 5% threshold for *α*-RRA algorithm. Genes were ranked according to the p-value and for genes with same p-value according to their mean LFC overall gRNAs.
- MAGeCK analysis was performed using the RRA algorithm, without normalization to controls. Genes were ranked according to the rank provided by MAGeCK.
- BAGEL analysis was performed only on experimental data because the algorithms needs a list of essential and non-essential genes as training sets. Creating this type of list on a set of simulated data would be arbitrary, since its quality cannot be compared to the currently available lists of essential and non-essential genes. BAGEL analysis was run without removal of low counts. Ranking of genes was performed based on Bayes factors.
- screenBeam analysis was performed without removal of low counts. Genes were ranked according to p-values.

## List of abbreviations

LFC: logarithmic fold change (base 2)
gRNA: guide RNA
RNA: ribonucleic acid
CRISPR: clustered regularly interspaced short palindromic repeats
Cas9: CRISPR associated 9
DNA: deoxyribonucleic acid
PCR: polymerase chain reaction
GO: gene ontology
RRA: robust rank aggregation
MOI: multiplicity of infection

## Declarations

### Availability of data and material

The simulation is available at https://github.com/imkeller/simulate_pooledscreen

`gscreend` (submission to Bioconductor pending) is available at https://github.com/imkeller/gscreend.

### Authors’ contributions

KI implemented the model and performed the analyses. GA performed the screening experiment. MB and WH designed and supervized the research. KI and WH wrote the paper.

## Acknowledgements

We would like to thank Sabine Ottinger for help with library preparation and acknowledge the support provided by the Genomics Core Facility of the European Molecular Biology Laboratory (Heidelberg, Germany).

## Supplement

**Figure S1:**
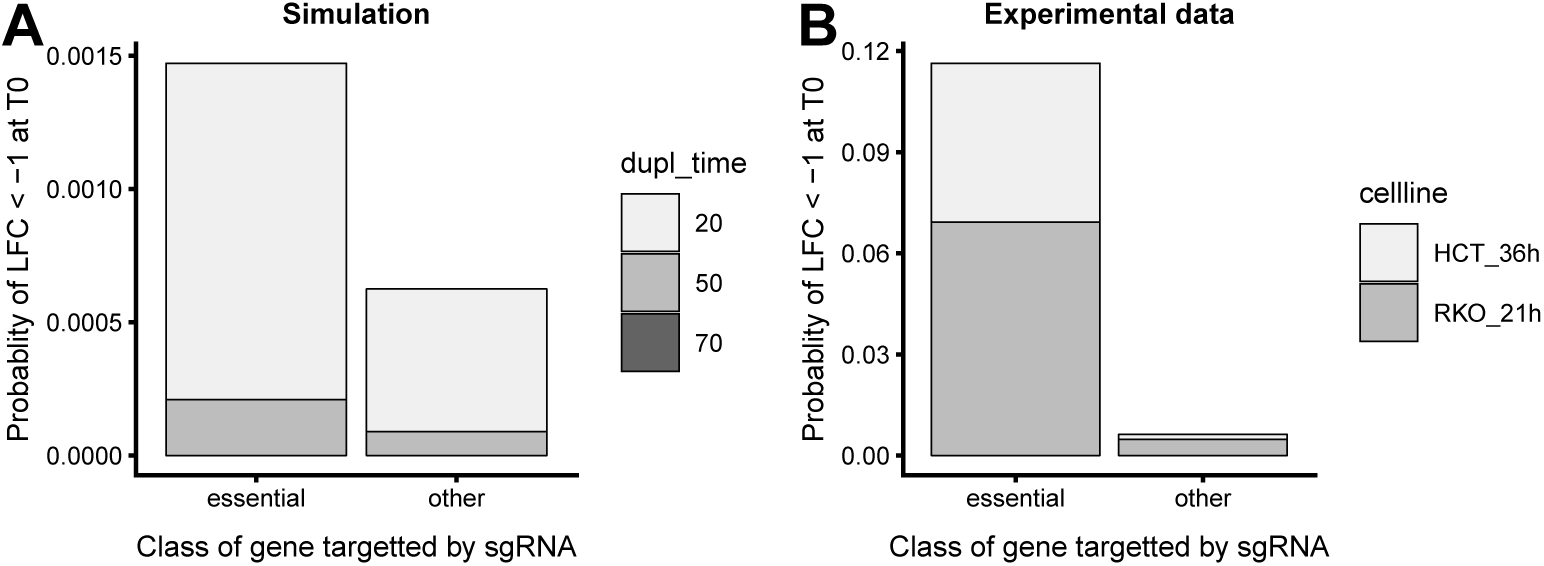
Fraction of gRNAs with reduced abundance at T0. The class of gene targeted by each category of gRNAs is indicated. Essential genes in (A) according to Hart et al.^38^. In (B) essential genes are all those that belonged to the set of growth rate reducing genes. A: Simulation with doubling times of 20, 50 and 70 hours. B: Experimental data from RKO (doubling time 21h) and HCT116 cells (doubling time 36h).

**Figure S2:**
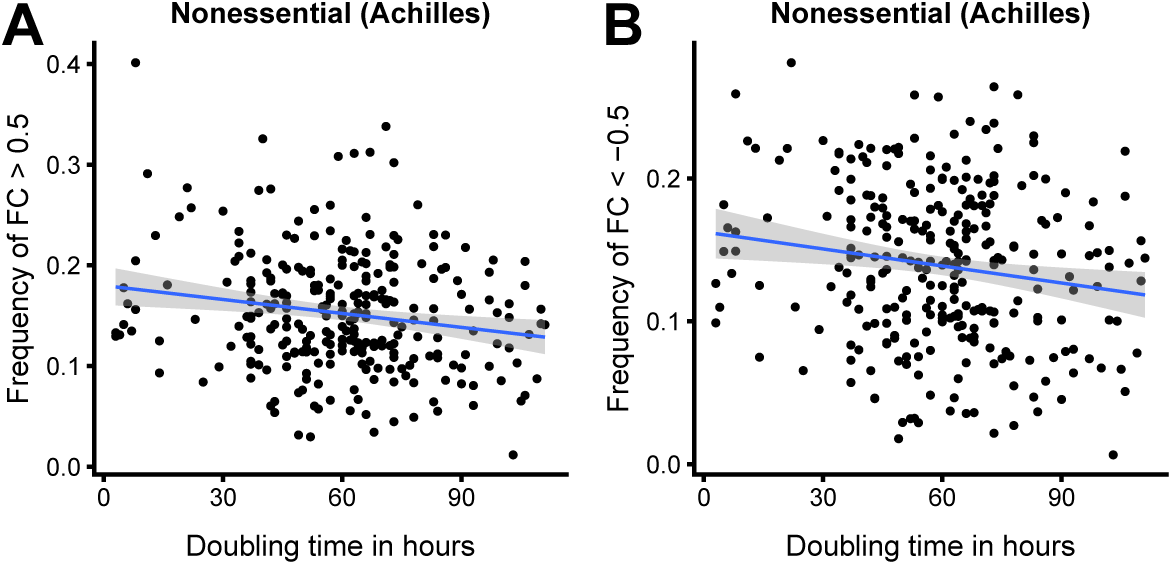
Screening data from the DepMap project shows higher frequency of LFC at higher cell growth rate. Analyses were performed only for gRNAs targeting non-essential genes. A: Positive LFC>+0.5. B: Negative LFC<-0.5.^3,39,40^

**Figure S3:**
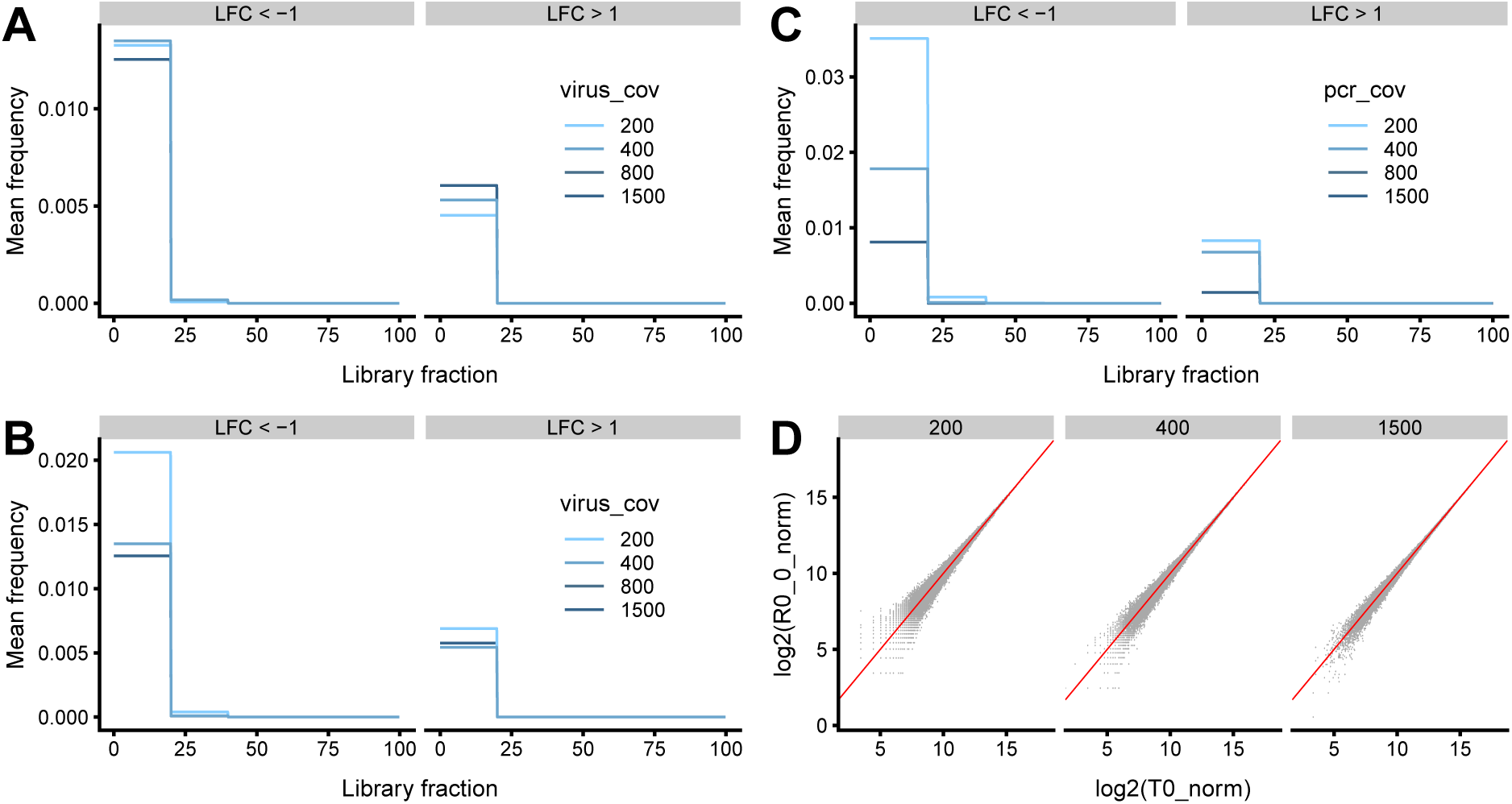
Effect of changing transduction or PCR coverage is small compared to the effects described in Figure 2. A: Frequency of extreme LFC between library sequencing and T1 for different values of transduction coverage. B: Frequency of extreme LFC between T0 and T1 for different values of transduction coverage. C: Frequency of extreme LFC between T0 and T1 for different values of PCR coverage. D: log2 gRNA counts at T1 vs. T0 for different values of PCR coverage.

**Figure S4:**
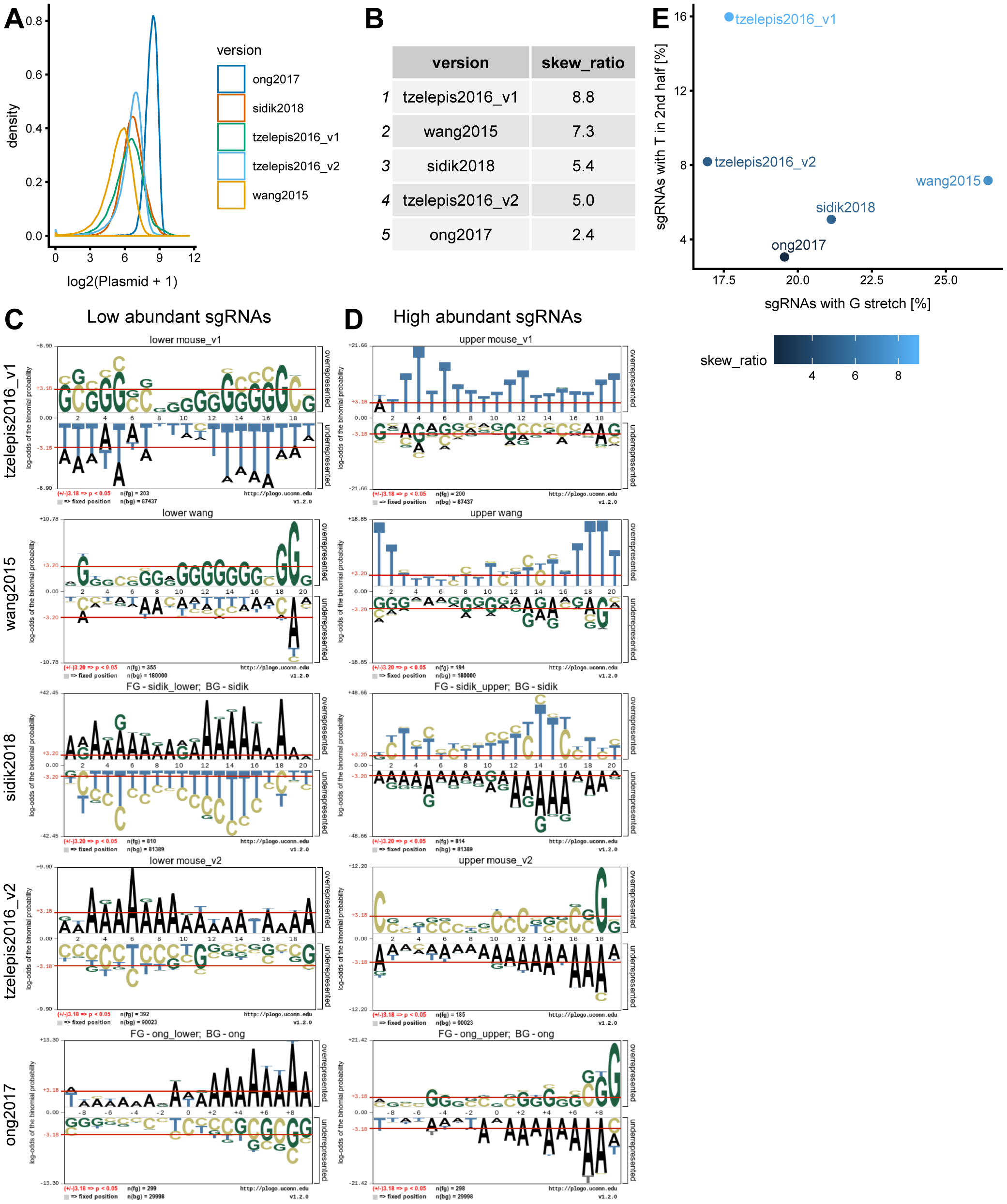
gRNA sequence features influence homogeneity of library distribution. A+B: Selection of five different CRISPR-KO libraries with different log2 gRNA count distributions (A, normalized to total counts) and library distribution widths (B)^1,27,41,42^. C+D: Probability sequence motifs for low (B) and high (C) abundant gRNAs in the libraries (ordered by decreasing distribution width). E: Percentage of gRNAs with more than three Ts in the 2nd half of the sequence (y-axis) and percentage of gRNAs with at least three consecutive Gs (x-axis).

